# Phagocytosis underpins the biotrophic lifestyle of intracellular parasites in the class Phytomyxea (Rhizaria)

**DOI:** 10.1101/2022.09.07.506935

**Authors:** Andrea Garvetto, Pedro Murúa, Martin Kirchmair, Willibald Salvenmoser, Michaela Hittorf, Stefan Ciaghi, Srilakshmy L. Harikrishnan, Claire M.M. Gachon, John A. Burns, Sigrid Neuhauser

## Abstract

Phagocytosis is a complex multi-gene trait of eukaryotes and allegedly one of the very defining features of this group. Although well documented for free-living unicellular eukaryotes and in specific cellular types of animals, data on phagocytosis in intracellular biotrophic parasites are scant. Indeed, the definition of intracellular biotrophy as complete reliance of a parasite on a living host, with which it constantly negotiates for the exchange of nutrients, is at odd with the consumption of particulate matter suggested by phagocytosis. Phytomyxea are intracellular biotrophic parasites infecting a broad group of hosts, ranging from plants to stramenopiles. They belong to the clade Rhizaria, where phagotrophy (i.e., phagocytosis as main mode to acquire nutrients) is the main mode of nutrition. The exact mode of nutrition of the biotrophic phytomyxea, including the agriculturally impactful phytomyxid *Plasmodiophora brassicae*, is still unresolved; despite investigations and the availability of molecular data. For other Phytomyxea, observations are patchy and molecular data altogether lacking. Here, using available genomic and transcriptomic data for Phytomyxea and the *de novo* sequenced transcriptome of the brown algae parasite *Maullinia ectocarpii*, we investigate the likelihood that the genetic machinery underpinning phagotrophy is conserved within the clade. We further document intracellular phagocytosis in *P. brassicae* and *M. ectocarpii* by transmission electron microscopy and fluorescent in situ hybridization. Our investigations confirm that molecular signatures underpinning phagocytosis exist in Phytomyxea and hint at a smaller subset of genes used for intracellular phagocytosis, which is similar between the two parasites. Microscopic evidence confirms the existence of intracellular phagocytosis, which seems to coexist with the manipulation of host physiology typical of biotrophic interactions. In both phytomyxid parasites investigated intracellular phagocytosis has adapted to the intracellular environment and seemingly targets specific organelles. Our findings shed light on the feeding behaviour of Phytomyxea, providing new molecular data for the class; and suggest a paramount and previously unrecognised role for phagocytosis in biotrophic interactions between host and parasite.

## Introduction

Often seen as a conserved and nearly universal trait in all major eukaryote lineages, phagocytosis underpins defining eukaryotic features such as the origin of endosymbiotic organelles and of the endomembrane system (Raven *et al.*, 2009; Yutin *et al.*, 2009). Phagocytosis is defined as the interiorization and internal digestion of particles larger than 0,5 μm (Flannagan *et al.*, 2012) and it is assumed to be one of the principal mode of nutrition in the majority of free-living heterotrophic microbial eukaryotes (thereby called phagotrophy), with the remaining feeding by osmotrophy (i.e. extracellularly digestion and/or absorption of molecules via the cell membrane).

Despite its pervasiveness among eukaryotes, we owe most of the information on this process to a special group of “professional phagocytes” from the immune system of vertebrate model organisms (Uribe-Querol & Rosales, 2020), for which molecular tools and laboratory experiments are possible. The investigation of phagotrophy and other trophic modes can be challenging in microbial eukaryotes and it relies heavily on microscopic observations and on the labelling/tracking of food items (Keymer *et al.*, 2017; Miura *et al.*, 2017; Godrijan *et al.*, 2022). Even then, the robustness of observational evidence may alone be insufficient in ascertaining the trophic niche occupied by an organism, and sometimes misleading (Not et al., 2007; Moreira & Lopez-Garcia, 2014). Indeed, phagotrophy is often used as an example of a range of “nearly behavioural” traits of microbial eukaryotes, the study of which requires a combination of molecular and laboratory-based investigations (Keeling, 2019).

Intracellular eukaryotic parasites can obtain macromolecules from their host via endocytosis, i.e., phagocytosis of solid food particles and pinocytosis of fluids and the solutes therein. For example, Apicomplexa such as *Plasmodium* spp. (Abu Bakar et al., 2010; Matz et al., 2020) and *Toxoplasma gondii* (Dou et al., 2014) ingest and digest macromolecules and pieces of host cell cytoplasm via endocytosis. The kinetoplastid *Trypanosoma cruzii*, has been reported to phagocytotically take up nutrients via the cytostome, a well-defined groove-shaped structure conserved from its free-living ancestors (Chasen *et al.*, 2020). Some intracellular parasites of fungi, oomycetes, and green algae (*Rozella polyphagi* and *R. allomycis*; Fungi, Cryptomycota) have been observed to actively engulf host cytoplasm and organelles, but also to recruit host mitochondria around their thallus, seemingly compensating for their own unstructured and depauperated ones (James *et al.*, 2013; Powell *et al.*, 2017). These findings place *Rozella* (together with the above-cited intracellular parasites of animals) in a particular trophic niche where conserved traits from free-living ancestors (e.g., phagotrophy) and derived traits co-evolved with the host (e.g., host manipulation) coexist within the same biotrophic organism.

With the notable exception of the photosynthetic chlorarachnids, phagotrophy is assumed to be the main mode of nutrition in almost all the free-living Rhizarians (Cavalier-Smith *et al.*, 2018). Within this clade, Phytomyxea (SAR, Rhizaria) are a class of unicellular eukaryotic parasites living as intracellular obligate biotrophs in plants and stramenopiles in marine, freshwater and terrestrial habitats (Bulman & Neuhauser, 2017; Cavalier-Smith *et al.*, 2018). The class is currently split into three main clades: the orders Plasmodiophorida and Phagomyxida (Hittorf *et al.*, 2020) and the recently described genus *Marinomyxa* (Kolátková *et al.*, 2020). Phylogenetically, Phytomyxea are sister to the free-living Vampyrellida (Sierra *et al.*, 2016; Cavalier-Smith *et al.*, 2018) and Aquavolonida (Bass *et al.*, 2018). Aquavolonida are a group of small, unicellular, free-living phagotrophic flagellates (Bass *et al.*, 2018). Vampyrellida are amoebae with different modes of prey item consumption, ranging from classic phagocytic predation to specialized protoplast feeding, where the prey cell wall is perforated and the amoeba enters the cell and phagocytise it from within (Hess & Suthaus, 2022). Phytomyxea use a very similar strategy to gain access to the host cell, piercing the cell wall with a sophisticated extrusome called Rohr and Stachel (Keskin & Fuchs, 1969; Aist & Williams, 1971). Distinctively, Phytomyxea reach the host cell as flagellated zoospores and penetrate into it as small unicellular protoplasts, later developing into larger intracellular multinucleate feeding plasmodia. Plasmodia can be of two types: short-lived (^~^ 7 days) sporangial plasmodia, developing into clusters of sporangia (i.e., sporangiosori) and directly releasing infective flagellated zoospores; or sporogenic plasmodia (i.e., sporosori), actively growing as biotrophs inside the living host cell (^~^ 3-4 weeks) before developing thick-walled overwintering resting spores. During that time sporogenic plasmodia induce hypertrophy of the infected cells which, coupled with induced hyperplasia of the tissue, leads to the formation of galls in the host (Murúa *et al.*, 2017; Olszak *et al.*, 2019). Manipulation of brassicaceae hosts by *P. brassicae* induces hypertrophied infected cells to act as physiological sinks, driving photosynthates from the aerial parts of the plant (Malinowski *et al.*, 2019) and inducing their accumulation as starch grains in the root (Ma *et al.*, 2022).

How Phytomyxea feed on their host has never been clearly elucidated and even the trophic mode of the model phytomyxean *P. brassicae* is still debated (Bulman & Neuhauser, 2017). Among the Phagomyxida, the diatom parasites *Phagomyxa* spp. have been observed to ingest the cytoplasm and organelles from their hosts by phagocytosis and accumulate the digested material in pigmented digestive vacuoles (Schnepf, 1994; Schnepf & Bulman, 2000). On the other hand, the lack of a conspicuous digestive vacuole and failure to detect engulfed host organelles has led to conclude that the brown seaweed-infecting phagomyxid *Maullinia ectocarpii* feeds by osmotrophy (Maier *et al.*,2000). Within the Plasmodiophorida, intracellular phagotrophy has been observed in the oomycete-infecting species *Woronina pythii* (Dylewski *et al.*, 1978) and *Octomyxa brevilegniae* (Couch *et al.*,1939; Pendergrass, 1950). Food vacuoles containing residues of cytoplasm and organelles from the host plant *Nasturtium officinale* (watercress) have also been found in *Hillenburgia nasturtii* (formerly *Spongospora subterranea* f. sp. *nasturtii;* Clay & Walsh, 1997; Hittorf *et al.*, 2020). Despite iconographic evidence supporting the existence of phagotrophy in *Plasmodiophora brassicae* (Williams & McNabola, 1967; Buczacki, 1983), a clear consensus on whether nutrition is dominated by osmotrophy, phagotrophy or consists of a mix of the two has not yet been reached (Dylewski, 1990).

Molecularly, complex and “behavioural” traits such as feeding modes are inherently difficult to investigate, since they are the final phenotypic outcome of a cohort of finely tuned genes involved in a range of overlapping (and often widely conserved) biological processes (Keeling, 2019). *In silico* predictions based on presence or absence of genome-wide molecular signatures identified in organisms known to possess a certain phenotypic trait can be used to infer the likelihood of the existence of that specific trait in other organisms, based on their genomic information (Burns *et al.*,2018). Direct observation, lab-based experiments and analysis of molecular data are complementary and have been successfully used to identify or rule-out phagotrophy in different groups of prasinophytes green algae (Bock *et al.*, 2021; Jimenez *et al.*, 2021).

In this study we used genomic and transcriptomic data from the plasmodiophorids *Plasmodiophora brassicae* and *Spongospora subterranea* (Schwelm *et al.*, 2015; Rolfe *et al.*, 2016; Ciaghi *et al.*, 2018a,b); and sequenced the transcriptome of the infective stage of the phagomyxid *Maullinia ectocarpii* to detect molecular signatures of phagotrophic behaviour (i.e. protein families present in well-known phagocytes) in the class Phytomyxea. We complemented results from these analyses with fluorescent and electron microscopy observations, to investigate if: (1) intracellular plasmodia engulf organelles and parts of the host cell, (2) the molecular machinery underpinning the phagocytic behaviour is present; (3) intracellular plasmodia express core genes involved in phagocytosis, similarly to other intracellular phagocytes (e.g., *Rozella allomycis*).

## Material and methods

### *M. ectocarpii* transcriptome: biological material, RNA extraction, sequencing and data processing

The model brown alga *Ectocarpus siliculosus* strain Ec32m (CCAP 1310/4) was used as a host for the co-cultivation of *Maullinia ectocarpii* (CCAP 1538/1) for RNA extractions. The pathosystem was maintained in half strength Provasoli medium at 15 °C, with a 12:12 h photoperiod, and an irradiance of 10 μE m^-2^ s^−1^. Quadruplicates of *E. siliculosus* Ec32m infected with *M. ectocarpii* were generated, harvested after 21 days with a 70 μm cell strainer (VWR, USA), and transferred immediately to ice-cold RNAlater (Ambion, Austin, TX, USA), stored overnight at 4 °C, and transferred at −80 °C until used for RNA extraction. Samples in RNAlater were thawed on ice, vortexed, and briefly spun down. 500 μL were transferred onto a pre-mixed Bead-matrix (Biozym D1034-MX). Samples were then spun down at 10000 g, 4 °C for 10 min, and RNAlater was carefully removed. Samples were immediately snap frozen in liquid nitrogen. Frozen material was subsequently homogenized with a FastPrep (MP Biomedicals, Santa Ana, CA, USA) for 40 s at 6 m s^-1^. This step was repeated three times and samples were returned into liquid nitrogen in between the three cycles to aid homogenisation and avoid RNA degradation. After the last homogenisation round, samples were transferred into liquid nitrogen and placed on ice. 450 μL buffer RLT (+ß-mercaptoethanol) from the Qiagen RNeasy Plant Mini Kit (Qiagen, Hilden, Germany) were added, samples were vortexed for 30 s and spun down briefly before processing them according to the manufacturer’s instructions with an additional ethanol (95%) washing step before RNA elution. RNA quality was tested on an Agilent Bioanalyzer 2100 (Agilent Technologies, Palo Alto, CA, USA). Poly-A selected strand specific library construction and paired-end sequencing (2x 125 bp on a HiSeq 2500 using v4 chemistry; Illumina, San Diego, CA, USA) was performed at the VBCF NGS Unit (Vienna, Austria). Quality of the raw reads was checked using FastQC v0.9.1 (Andrews, 2010). Illumina adapters were removed and only good quality reads (sliding window 5 bp; average quality score > 20) with a minimum length of 50 bp were kept using Trimmomatic v0.36 (Bolger *et al.*, 2014). Bacterial contamination was removed from the remaining reads using DeconSeq v0.4.3 (Schmieder & Edwards, 2011). Reads from the mock and infected samples were separately mapped against the Ec32m reference genome v2 (Cock *et al.*, 2010) using Bowtie2 v2.2.4 (Langmead & Salzberg, 2012). Unmapped reads from the mock samples were *de novo* assembled into transcripts using Trinity v2.4.0 (Grabherr *et al.*, 2011) with default settings for k-mer size (25 bp) and minimum contig length (200 bp). These transcripts were further used as a reference to filter out host reads from the infected samples and select only reads unambiguously assigned to *M. ectocarpii* (i.e. unmapped reads of this filtering step). Remaining reads were *de novo* assembled into transcripts using Trinity with default settings, thus constituting *M. ectocarpii* transcriptome. Read counts (i.e. gene expression) of the assembled transcripts was estimated using RSEM (Li & Dewey, 2011) included in the Trinity suite. Only transcripts with FPKM (fragments per kilobase per million reads) values greater than one were kept for downstream analysis. Completeness of the transcriptome was verified using BUSCO v5.2.2 running in transcriptome mode with the eukaryote_odb10.2019-11-20 reference gene set (Simão *et al.*, 2015). *Maullinia* proteome was inferred using the longest open reading frames and the protein coding genes predicted by Transdecoder v5.0.2 (https://github.com/TransDecoder) with default settings and used in downstream analyses. Functional annotation of the predicted genes was achieved using InterProscan v5 (Jones *et al.*, 2014).

### Additional molecular data

Transcriptome data from an Austrian population of *Plasmodiophora brassicae* were taken from Ciaghi *et al.*, 2018b. Publicly available genomic data were taken from *P. brassicae* strains e3 (Schwelm *et al.*,2015) and PT3 (Rolfe *et al.*, 2016); and *Spongospora subterranea* strain K13 (Ciaghi *et al.*, 2018a).

### *In silico* predictions of trophic mode

BUSCO v5.2.2 was run in proteome mode against the eukaryote_odb10.2019-11-20 reference gene set (Simão *et al.*, 2015) to assess completeness of all inferred proteomes, allowing for accurate predictions of trophic modes (Liu *et al.*, 2021). Genomic and transcriptomic data from all three species of phytomyxean parasites in this study show a high degree of BUSCO completeness (< 105 missing BUSCOs over the total 255 BUSCOs in the eukaryota_odb10 database), indicating that their trophic mode can be accurately assigned by TrophicModePredictionTool (Supplementary Figure 1).

The TrophicModePredictionTool tool (Burns *et al.*, 2018) was used to predict the trophic mode of the investigated organisms *in silico*, based on the molecular signatures for phagocytic, photosynthetic, and prototrophic capabilities (i.e. organisms capable of synthetising arginine, lysine, threonine, biotin, vitamin B1, B2 and B6). The code (available at https://github.com/burnsajohn/predictTrophicMode) was run in the default mode. Prediction scores enumerate the probability that an organism has the genetic toolbox to carry out the indicated function on a scale of 0 to 1. A probability above 0.5 suggests that an organism has the capacity to utilise the indicated function, higher scores indicate a higher degree of confidence.

Besides the three main trophic modes listed above, special form of phagocytosis such as that of the extracellular parasite *Entamoeba histolytica* and of the intracellular parasite *Rozella allomycis* are predicted via an emended subset of molecular signatures of phagocytosis. Predictions were visualized as bar charts and by projecting the 4-dimensional probability values onto a 3D tetrahedral shape representing the three trophic modes (or their absence) using scripts modified from the R package “pavo”(Doucet *et al.*, 2013). For static visualization of the trophic mode of an organism, the 3D tetrahedral shape with the summary prediction from each organism plotted onto it is finally rendered as a 2D circular Mollweide projection as described in Bock *et al.*, 2021 and Jimenez *et al.*, 2021. A detailed overview of the genes best matching the predictive molecular signatures are presented in Supplementary Material 1 for the comparison between *P.brassicae* e3 genome and *P. brassicae* transcriptome; as well as for the comparison between *P. brassicae* transcriptome and *M. ectocarpii* transcriptome.

### Fluorescent in situ hybridisation and optical microscopy

*P. brassicae* was grown on the host plant *Brassica rapa* var. *pekinensis* (cultivar “Granat”) for 61 days before collection of root galls, thus allowing for the presence of a high number of plasmodia at different stages of development. Plants were grown at 20°C with a 12:12 photoperiod and an average irradiance of 135 μE m^-2^ s^−1^. Galls were thoroughly rinsed in tap water to remove soil residues and preserved in Histofix 4% (phosphate-buffered formaldehyde solution, Carl Roth, Germany) for ^~^1 hour. Following fixation, galls were dehydrated in ascending ethanol series: 10 min in 50% ethanol, twice 10 min in 70% ethanol and final storage in absolute ethanol. Galls were prepared for FISH staining following the procedure detailed in Schwelm et al., 2016, with few modifications. Briefly, galls were hand cut into thin sections and rinsed for 10 min in hybridization buffer (900 mM NaCl, 20 mM Tris HCl pH 7.5, 35% formamide, 0.01% SDS) before incubation overnight at 46°C in the dark in hybridization buffer, amended with 50 ng of the FISH probe Pl_LSU_2313 (Table 1). Samples were washed twice for 20 min in washing buffer (900 mM NaCl, 20 mM Tris HCl pH 7.5, 5 mM of NaEDTA pH8, 0.01% SDS) at 48°C. Samples were then incubated for 20 min in Hoechst 33342 (Thermo Scientific, USA) diluted 1000X in distilled water, before being mounted in Vectashield (H-1000, Vector Laboratories, USA). *M. ectocarpii* was grown on *Ectocarpus siliculosus* Ec32m male gametophyte or *Macrocystis pyrifera* CCAP1323/1 female gametophyte (same culture conditions specified above) for one month before collection. Fixation and FISH staining was achieved in the same way described for *P. brassicae* with the following adjustments. After fixation in 4% Histofix infected algae were incubated for 2 min in 30% H2O2 to increase cell wall permeability and then dehydrated in ascending ethanol series. The hybridization was performed at 46°C overnight in the dark in hybridization buffer amended with 50 ng of probe MauJ17 (Table 1). Slides were observed with a Nikon Eclipse Ti2-E microscope equipped with an Andor Zyla 5.5sCMOS monochrome camera and Nikon CFI Plan-Fluor 40×/0.75 NA and 60×/0.85 NA objectives. The excitation wavelength for Hoechst 33342 was 365 nm, whereas it was 490 nm for FISH probes (Table 1). The NIS Elements software (Nikon, Japan) was used for image analysis and post-processing (generation of overlaid images, z-stack analysis and export of z-stack as videos). Final figures were composed using Inkscape 0.92.4 (Inkscape Project).

**Table 1:**
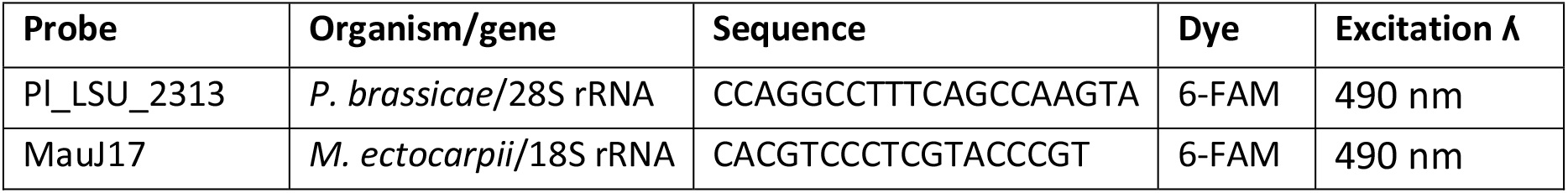
Fluorescent In Situ Hybridization probes used in this study.

### Transmission electron microscopy

For TEM, *M. ectocarpii* was grown on healthy female gametophytes of *Macrocystis pyrifera* CCAP 1323/1 in 1/2 strength Provasoli medium, at 10°C, under 2–6 μE m^-2^ s^-1^ white light irradiation and 12:12 h photoperiod. Biological material was chemically fixed and processed as per Murúa *et al.* (2017). Briefly, the biomass was immersed in a solution composed of 2.5% glutaraldehyde, 0.1 M cacodylate buffer at pH 7.4, 0.5% caffeine, 0.1% CaCl_2_ and 0.3% NaCl in Provasoli-enriched seawater (PES) for 2-3 days. Post fixation staining was achieved with 1% OsO_4_ and 2% uranyl acetate. After dehydration in acetone series, samples were embedded in Spurr resin and polymerised at 60-70°C. Blocks were cut using a Leica UC6 ultramicrotome and counterstained on copper grids with lead citrate. Imaging was achieved with a JEM-1400 Plus (Jeol, Akishima, Tokyo, Japan) TEM with an AMT UltraVue camera (Woburn, MA, USA). For TEM imaging of *P. brassicae*, root galls of *Brassica rapa* var. *pekinensis* were collected from field material in Weer (Tirol, Austria) in September 2018. Specimens were chemically fixed with 2.5% glutaraldehyde in 0.1 M cacodylate buffer containing 10% sucrose for 1 h at 4 °C, rinsed with cacodylate buffer and post fixed with 1% osmium tetroxide in 0.05 M cacodylate buffer for 1 h at 4 °C. After washing in cacodylate buffer, samples were dehydrated with an increasing acetone series and embedded in EMbed 812 resin. Cross sections of root galls were cut with a diamond knife (Diatome, Switzerland) and an Ultracut UCT (Leica, Austria), mounted on grids, stained with lead citrate and examined with a Libra 120 energy filter transmission electron microscope (Zeiss, Germany). Images were made with a TRS 2 × 2k high speed camera (Tröndle, Germany) and an ImageSP software (Tröndle, Germany).

## Results

### *In silico* prediction of trophic modes of Phytomyxea using genomic and transcriptomic signatures

All analysed phytomyxids datasets bear molecular signatures of phagotrophy (Fig. 1, Supplementary Table 1). *Plasmodiophora brassicae* (e3 and PT3) and *S. subterranea* (SSUBK13) genomes score high for phago-prototrophy (red numbers 11, 12 and 18 on the Mollweide projection in Fig. 1). The prediction scores from the genome data in *P. brassicae* are approximatively 60% for prototrophy (e3 = 0.615 and PT3 = 0.612) and are similar for general phagotrophy (e3 = 0.700; PT3 = 0.600). The prototrophy score for *Spongospora subterranea* is lower (SSUBK13 = 0.500, bar chart in Fig. 1), as is the score for general phagotrophy (SSUBK13 = 0.552). When the subset of signatures predicting *Rozella*-like intracellular phagotrophy is considered the probability scores increase to nearly 100% for the genome datasets (e3 = 0.978; PT3 = 0.983 and SSUBK13 = 0.967; bar chart in Fig. 1). The probabilities for photosynthesis and entamoebid-like phagotrophy (a second peculiar mode of phagotrophy mostly observed in extracellular endoparasites such as *Entamoeba*) remain below the threshold of 50% in the genomic data (Supplementary Table 1). When the proteomes inferred from the transcriptomes of *P. brassicae* and *M. ectocarpii* are tested the score for general phagotrophy and prototrophic predictions are very low: in *P. brassicae* the prediction score for general phagotrophy decreases to 0.018 and prototrophy to 0.087, in *M. ectocarpii* phagotrophy scores 0.209 and prototrophy 0.179. The *Rozella-like* phagotrophy remains high with a score of 0.838 in *P. brassicae* and 0.894 in *M. ectocarpii* (bar chart in Fig. 1). Presenting these data as a Mollweide projection, the transcriptome datasets are placed with other “Parasite” mapping close to the intracellular fungal parasite *Rozella allomycis* (red numbers 9 and 13 and black number 16 in the Mollweide projection in Fig. 1), while the genomic datasets are in the phago-prototroph area. The assignment to the “Parasite” area in the Mollweide projection highlights a low score (<0,5) for the main trophic categories (i.e. general phagotrophy, prototrophy and photosynthesis), but does not exclude the assignment to specialised sub-categories of phagocytosis (i.e. *Entamoeba* or *Rozella-like* phagocytosis) as highlighted by the bar chart in Fig.1.

**Fig. 1.**
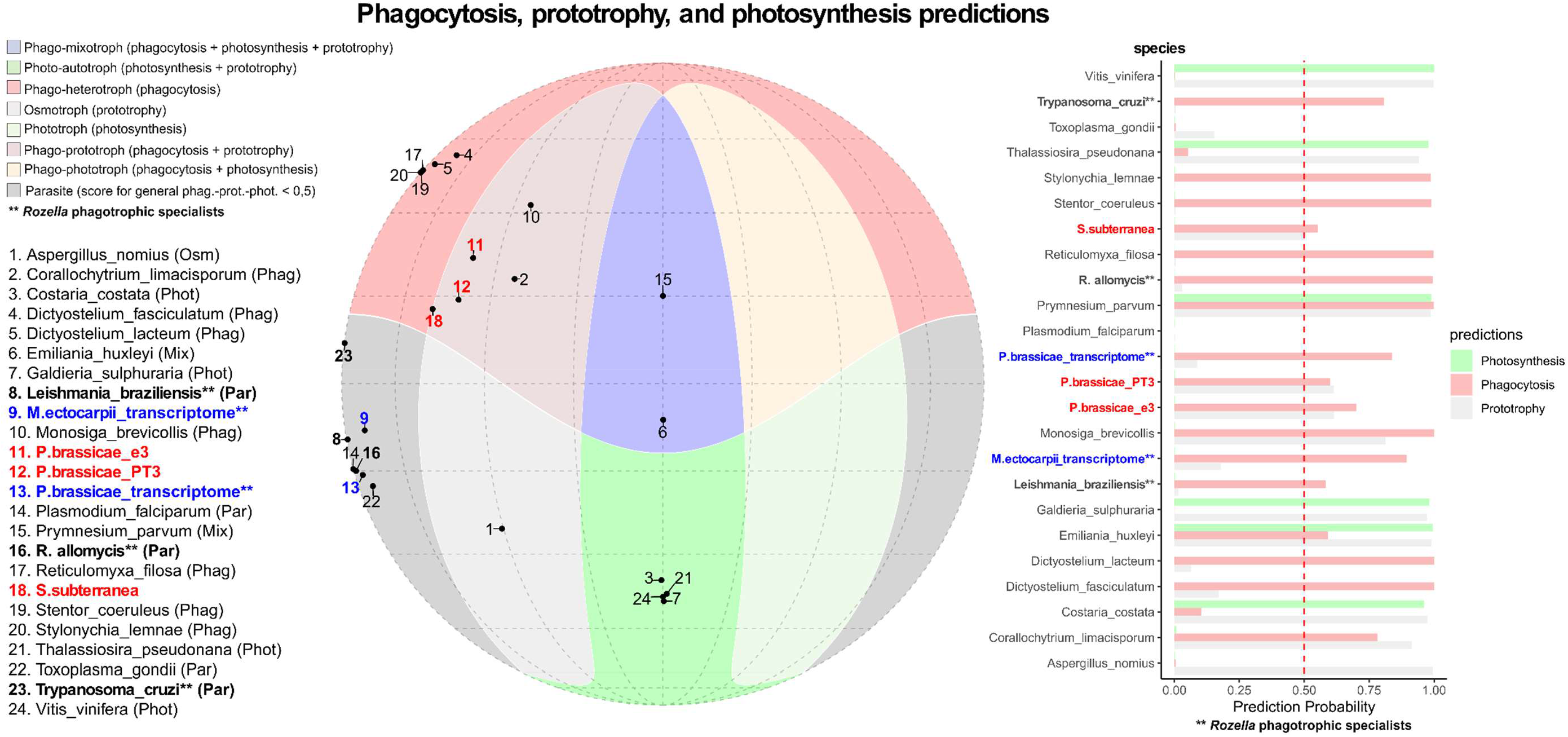
Mollweide projection showing the position of predicted phagotrophs, prototrophs, and photosynthetic organisms; and bar chart showing the scores of individual prediction probabilities for the same organisms. Colored regions (red, blue, beige, green) indicate overlapping areas where individual predictions were >0.50. Dark-gray shaded regions indicate areas where all three predictions were <0.50. Note that phagotrophic specialists (such as *Rozella*-like phagotrophs) do not map the three main trophic categories and fall in the grey area due to the Mollweide projection only using the general phagocytosis prediction for each organism. Each numbered black dot indicates one of the 19 organisms used as a reference to test the model. The same organisms are represented in the bar chart where prediction probability scores are shown as coloured bars (green=photosynthetic, grey=prototroph, brown=phagotroph). The 0.5 threshold, above which a prediction is deemed valid, is indicated by the red dashed line. Names in red/bold indicate the phytomyxid genomes tested in this study whilst blue/bold indicates phytomyxid transcriptomes; names in bold and followed by double asterisks (**) indicate organisms for which the strongest prediction is *Rozella*-type intracellular phagotrophy.

In *P. brassicae* (for which both genomic and transcriptomic data are available), a detailed look at the molecular signatures highlighted that nearly half (14/29) of the phagotrophy-related genes driving the genome apart from the transcriptome were associated with cilia/flagella (as per their GO term annotation, Supplementary Material 1). Within the predictive model, flagella and cilia are describers of the phago-prototrophic niche, which accommodates organisms using these structures to feed (e.g. Choanozoa such as *Monosiga brevicollis*; Fig. 1).

On the other hand, trophic predictions for the transcriptomic datasets of *P. brassicae* and *M. ectocarpii* were similar (Fig. 1). In *P. brassicae* genes associated with phagotrophic signatures in the transcriptome were linked to GO terms involving the cytoskeleton (14/40), the cytosol (6/40) and the mTOR complexes (5/40); including the GO terms TORC2 complex, Seh1-associated complex and the lysosome gene RRAGA (Supplementary Material 1). A closer look at the predicted functions highlights their potential involvement in processes such as signal transduction, cell reorganization/polarization, metabolism and cell cycle. In particular Ras GTPases, mTORC1 and mTORC2 complexes are strong descriptors of *Rozella*-like phagotrophic behaviour and describe nearly half (10/21) of the signatures shared between the transcriptomes of *P. brassicae* and *M. ectocarpii* (Supplementary Material 1).

### Microscopic evidence of phagocytosis in intracellular plasmodia of *P. brassicae* and *M. ectocarpii*

Microscopic observation of intracellular biotrophic plasmodia of *P. brassicae* and *M. ectocarpii* support phagotrophy of host organelles by the parasites. Mature feeding plasmodia of *P. brassicae* (Fig. 4, A) could be recognized by the high number of small nuclei (Fig. 4, B’-D’; small blue dots) in the absence of cytoplasm cleavage. The plant nucleus was still present and could be distinguished from the parasite nuclei by its larger dimensions (Fig. 4, B-B’; white triangle). Plasmodia filled up the host cells entirely (Fig. 4; A-D; green), leaving little free space within the cell wall. Abundant starch grains were easily identified in DIC microscopy by their shape, size, hyaline texture and tridimensional appearance (Fig.4; B”-D”). A high number of starch grains was located between the plant cell wall and the parasite plasma membrane, pressed against the plasmodium as if superficially “plugged” in membrane pockets (Fig. 4, B-B’ and D-D’). Many starch grains were also found to be completely enveloped by the parasite plasmodium, often contiguous to other engulfed and “plugged” starch grains, giving the plasmodium an overall “sponge-like” or “trabecular” aspect in fluorescence microscopy (Fig.4, B’-D’). Two starch grains were entirely surrounded by the plasmodium (Fig.4; C, C’ and C”, white arrowheads; videos in Supplementary Material 2) as highlighted by the presence of green hue around them and in the focal planes above and below them.

*E. siliculosus* cells infected by mature *M. ectocarpii* were easily distinguishable thanks to the clear signs of hypertrophy (Fig.4 E, white square). The plasmodium shown (Fig. 4, F-H) occupied the majority of the space within the host cell wall, as indicated by the green FISH staining of its cytoplasm. The plasmodium was multinucleated (Fig. 4; F-H and F’-H’; blue signal) and showed vacuolar structures where no green fluorescence could be observed (Fig. 4; F-H and F’-H’; white arrowheads and asterisk; videos in Supplementary Material 2). Some vacuoles contained refractive structures consistent with the phaeoplasts of *E. siliculosus* (Fig. 4; G, G’ and G”; white arrowheads) whilst other did not (Fig. 4; G, G’ and G”; white asterisk). Phaeoplasts were also observed to be “plugged” in membrane pockets (Fig. 4; F, F’, F” and H, H’, H”; white arrowhead), much like starch grains in *P. brassicae*. Scansions of the entire volume of the investigated plasmodia along the z-axis are available as videos (Supplementary Material 2); allowing for a better visualization of the host organelles engulfed by the parasites. To further strengthen our observations, we performed a FISH experiment on *M. ectocarpii* infecting the female gametophyte of the kelp *Macrocystsis pyrifera*. Even in this case, phagocytosis was observed as highlighted by the observation of phagocytic vacuoles and the late phagocytosis of the host nucleus (Supplementary Figure 2 and Supplementary Material 3).

### Ultrastructural evidence of phagocytosis in intracellular plasmodia of *P. brassicae* and *M. ectocarpii*

Plasmodia and thick-walled resting spores of *P. brassicae* were observed inside the cortical cells of *Brassica rapa* subs. *pekinensis.* Plasmodia can be discriminated from the plant host because of the high amount of lipid globules stored within the cytoplasm (absent from healthy plant cortical cells), the different electron density of the cytoplasm and the electron light mitochondria with sparse tubular cristae (Fig. 2). Multinucleate plasmodia occupy most of the host cell, leaving space only for the host nucleus, small vacuoles and few smaller organelles (like mitochondria) embedded in a film of plant cytoplasm appressed to the cell wall. Parasite nuclei were clearly distinguishable from the plant nuclei, because of their rounder shape and smaller size (Fig. 2, A-A’). The overall shape of the plasmodium was irregularly lobed, to the extent that often it was impossible to clarify whether a single highly lobed or many different plasmodia were inhabiting the same host cell. (Fig. 2, A-A’) Lobes of different shape and size were often found surrounding and/or closely appressed to starch grains, originated from desegregated amyloplasts (Fig. 2, B-B’; C-C’). Often plasmodial lobes seemed to encircle and close around starch grains (Fig. 2, C-C’) and in one occasion one of those granules was found to be completely surrounded by the plasmodium (Fig. 2, A, StG). The mitochondria of *P. brassicae* were found to be generally electron-translucent and contained less cristae than the lamellar plant mitochondria (Fig. 2, B-B’; C-C’. Mitochondria in thick-walled resting spores were much better defined in their ultrastructure and are overall more electron opaque (Supplementary Figure 4).

**Fig. 2:**
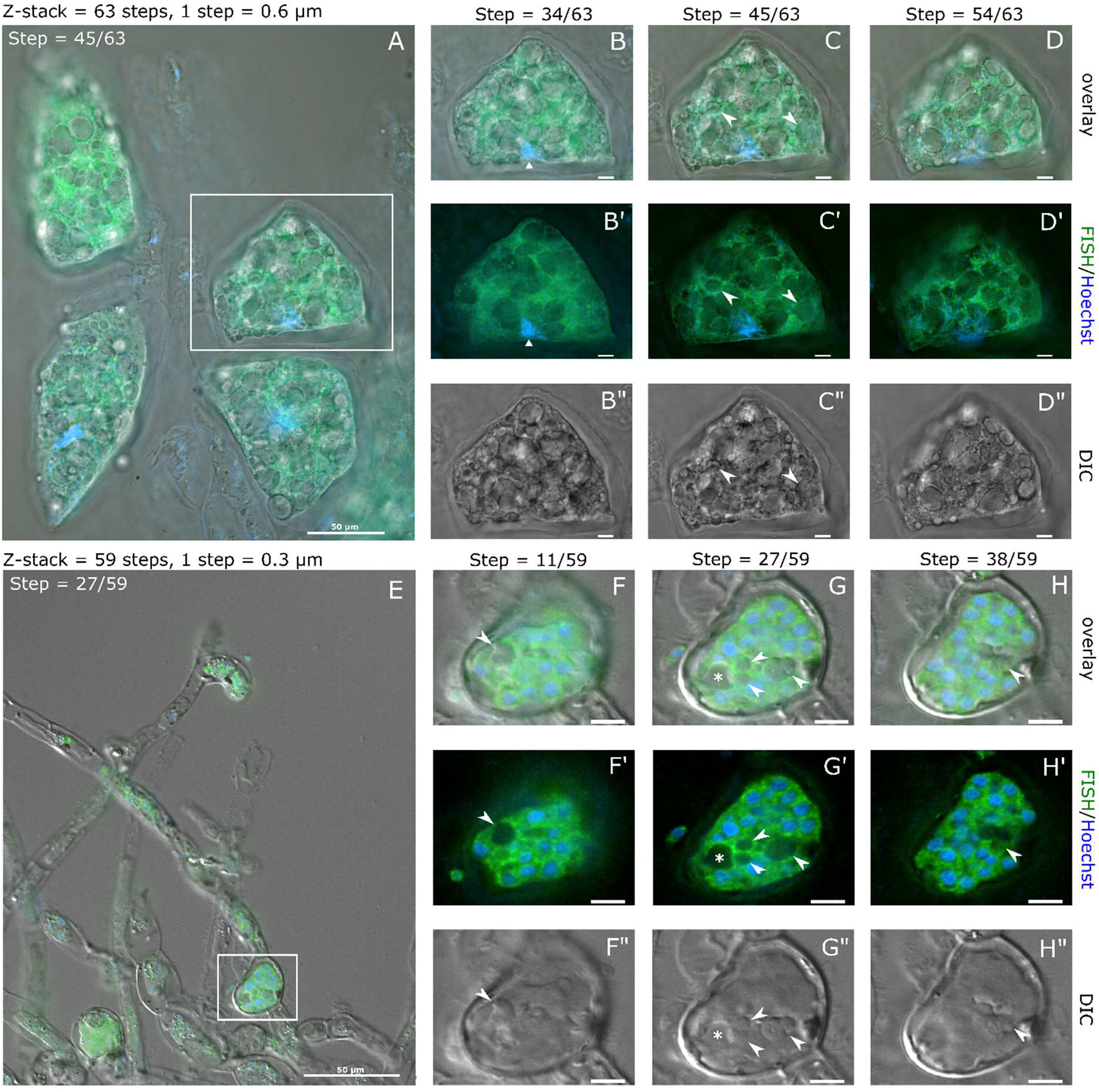
Plasmodia of *Plasmodiophora brassicae* growing within the cortical cells of the root of *Brassica rapa* subs. *pekinensis*. Annotations are provided in a separate image from the original picture (e.g. annotations for A in A’, etc.) **A-A’:** overview of the interface between the plasmodium and its host. Note the starch granule surrounded by the plasmodium. **B-B’:** Starch grains from desegregated amyloplasts are surrounded by plasmodial protrusions, some of which in closely appressed to the granule surfaces. **C-C’:** Detail of a plasmodial pseudopodium-like process protruding between two starch grains. **In all pictures** note the presence/absence of lipid droplets and the different electron opacity/organisation of the mitochondria, used as main distinctive features to tell apart host and parasite. List of abbreviations: **HN** = host nucleus, **Hn** = host nucleolus, **PN** = parasite nucleus, **Pn** = parasite nucleolus, **white dots** = parasite lipid droplets, **HMt** = host mitochondria, **asterisks** (**\***) = parasite mitochondria, **CW** = host cell wall, **StG** = starch grains, **black lines** indicate the plasma membrane of the plasmodium. Scale bars: A-A’ = 2500 nm, B-B’ = 500 nm, C-C’ = 1000 nm.

**Fig. 2:**
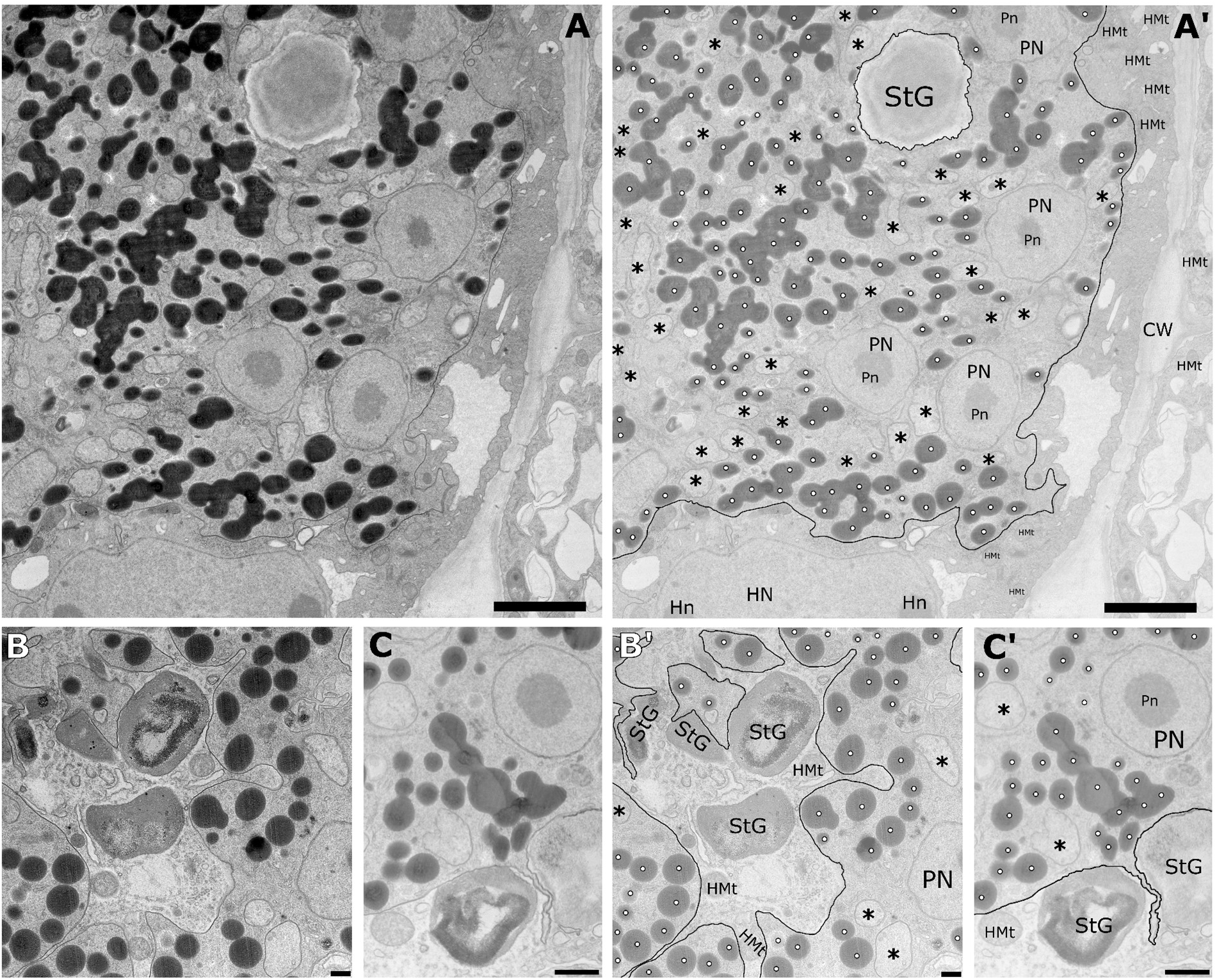
Optical and fluorescence micrographs provide evidence of the internalisation of host organelles in intracellular plasmodia of *Plasmodiophora brassicae* in *Brassica rapa* var. *pekinensis* (A-D) and of *Maullinia ectocarpii* in *Ectocarpus siliculosus* Ec32m (E-H). Images have been captured using differential interference contrast microscopy (DIC; B”-D” and F”-H”) and fluorescence microscopy (B’-D’ and F’-H’) and subsequently overlaid (A-D and E-H). FISH-probes specific to the 28S rRNA gene of *P. brassicae* and 18S rRNA gene of *M. ectocarpii* were used to highlight the ribosome rich cytoplasm of the parasites (green). Hoechst staining highlighted the nuclei of both parasite and host (blue) in fluorescence microscopy. White squares in A and E indicate the plasmodia shown in detail in B-D and F-H respectively. The white triangle in B points toward the Hoechst-stained host cell nucleus, whilst white arrowheads in C indicate two completely internalised starch granules. Arrowheads in F-H highlight engulfed algal phaeoplasts, whilst the asterisk in G indicate a vacuole. Focal planes represent a high (B, B’, B” and F, F’, F”), a central (C, C’, C” and G, G’, G”) and a low (D, D’, D” and H, H’, H”) layer from z-stacks containing entire plasmodia. In B to D, scale bars 10 μm; in F to H, 5 μm.

Feeding plasmodia of *M. ectocarpii* were observed in intercalary and tip cells of the filamentous female gametophytes of *Macrocystis pyrifera* (Fig. 3). Plasmodia readily occupy the whole host cell, initially taking up the space of the central vacuole, thereby pushing the organelles towards the periphery of the cell. *M. ectocarpii* plasmodia are easily discriminated from the host cell by the absence of phaeoplasts and because of the difference in the cytoplasmic electron-density (i.e., denser in the alga; Fig. 3). Electron dense mitochondria with tubular cristae have been noticed in the algal host. In the plasmodia of *M. ectocarpii* mitochondria are not as visible: putative mitochondria appear as double-membrane bounded electron translucent structures without clearly discernible tubular cristae (

**Fig. 3:**
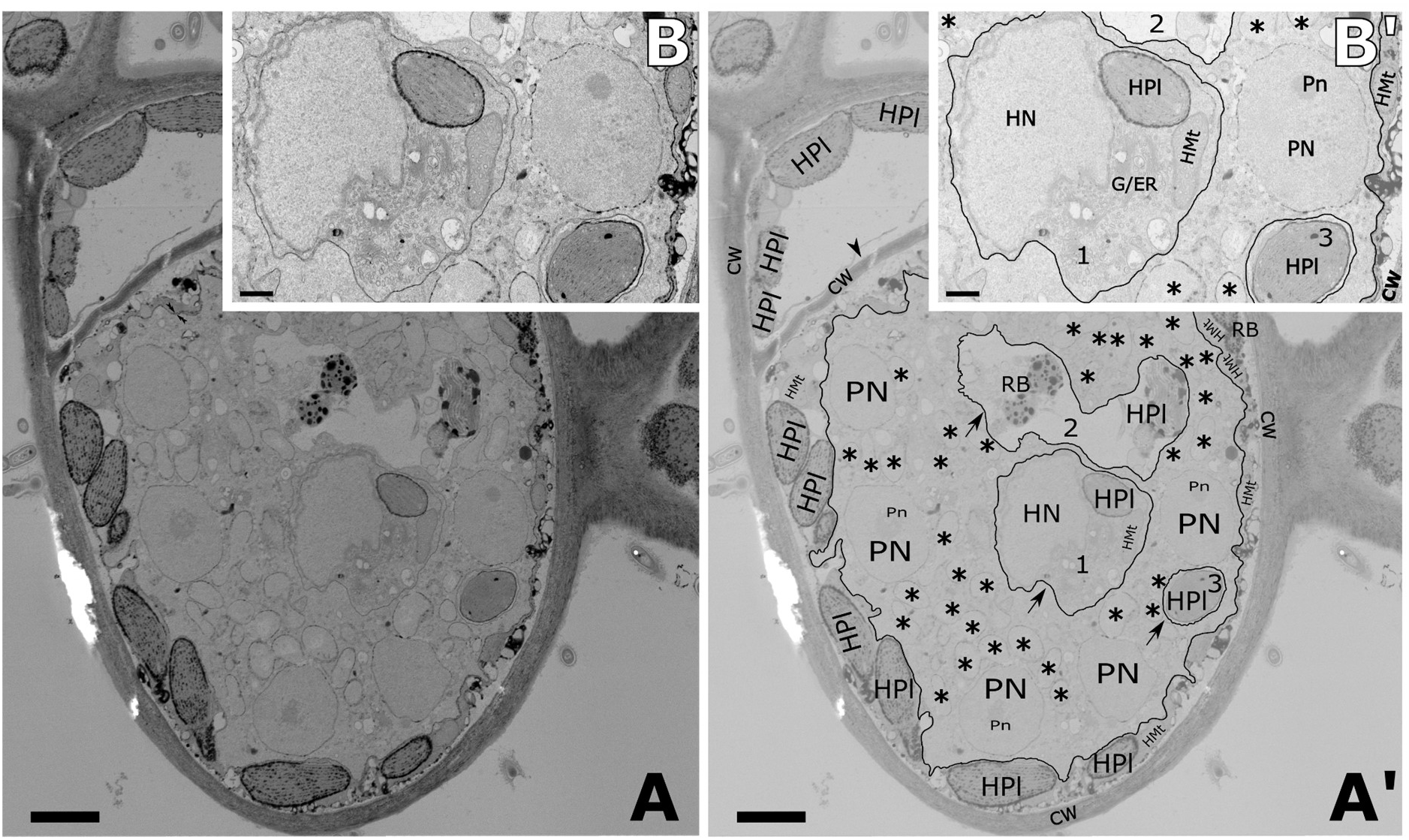
Plasmodium of *Maullinia ectocarpii* growing within a cell of the gametophyte of *Macrocystis pyrifera*. Annotations are provided in a separate image from the original picture (e.g. annotations for A in A’, etc.) **A-A’:** overview of the interface between the plasmodium and its host. Note the three vacuoles containing algal mitochondria, phaeoplasts and nucleus bound by membrane and surrounded by the parasite plasmodium (arrows). **B-B’:** Close up of A-A’ highlighting details of the algal organelles surrounded by the parasitic plasmodium. Note the difference in electron opacity/granularity between the parasite and host cytoplasm used as main distinctive features to tell apart host and parasite. Secondarily, note the absence of clear tubular cristae in the putative mitochondria of *M. ectocarpii*. List of abbreviations: **HN** = host nucleus, **Hn** = host nucleolus, **PN** = parasite nucleus, **Pn** = parasite nucleolus, **HPI** = host phaeoplasts, **HMt** = host mitochondria, **asterisks** (**\***) = potential parasite mitochondria, **CW** = host cell wall, **RB** = residual bodies, **black lines** indicate the plasma membrane of the plasmodium, **Arrows & Numbers** = potential vacuoles, **Arrowhead** = cell wall separating infected and non-infected sectors within the same cell. Scale bars: A-A’ = 2500 nm, B-B’ =500 nm, C-C’ = 1000 nm.

**Fig. 3.**
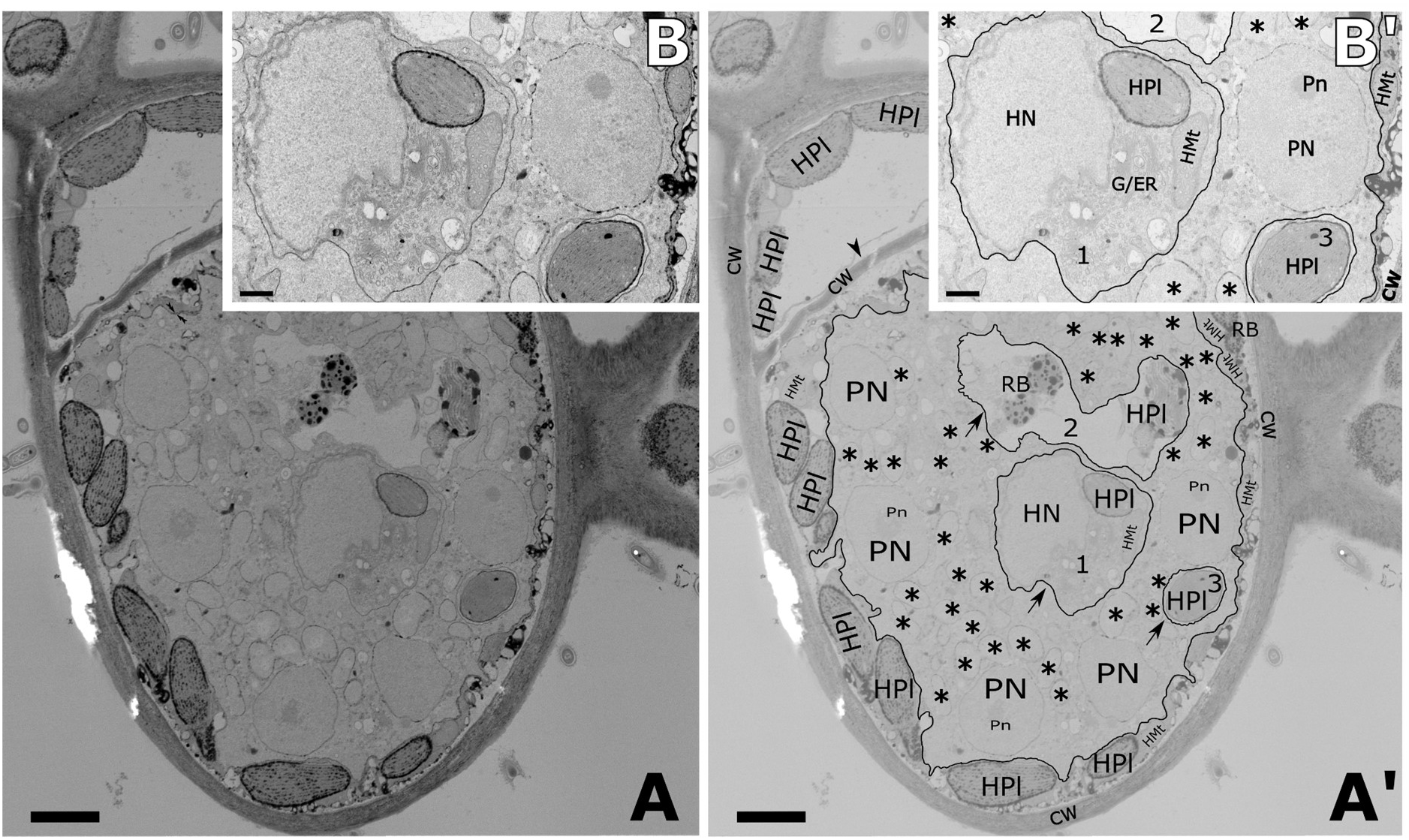
A-A’; B-B’). Comparably with observations in *P. brassicae*, in mature *M. ectocarpii* plasmodia/zoosporangia, mitochondria within zoospores show a higher level of structuration, being electron-denser and with well-organised tubular cristae (Supplementary Figure 4). Plasmodia are irregular and sometimes structures similar to pseudopodia can be observed, especially in very young, developing plasmodia which do not yet fill the host cell (Supplementary Figure 3). Vacuoles are often observed within *M. ectocarpii* plasmodia and differ in size and content (Fig. 3 A’-B’, arrows and numbers 1-3). Vacuoles can be nearly empty (electron translucent), but most vacuoles are either loosely filled with degraded material (Fig. 3 A’-B’, 2) or filled with host organelles and cytoplasm (Fig. 3 A’-B’, 1 and 3). In Fig.3 B-B’ vacuole number 3 can be observed containing a phaeoplast, with little to no space for other structures. A second, bigger vacuole (number 1) contains the host nucleus together with a phaeoplast, one host mitochondrion and host cytoplasm, in turn containing membranous structures interpreted as endoplasmic reticulum and/or Golgi apparatus. An even bigger vacuole (number 2) can be observed in Fig. 3, A-A’, within which a clearly degraded phaeoplast and two residual bodies, potentially representing a further stage in paheoplast degradation, can be observed. The presence of degraded phaeoplasts in vacuole 2 suggests that these have been isolated from the rest of the host cytosol and digested. Presumably, vacuoles 1 and 3 are bound to undergo the same process. The plasmodium itself is multinucleate but it has not yet undergone cytodieresis and zoospore cleavage.

## Discussion

In this study, by analyzing complementary lines of evidence, we demonstrate that phagocytosis is a trait that phytomyxean parasites conserved from free-living Rhizarian ancestors, adapting it to the intracellular environment where it underpins the biotrophic interaction and where it coexists with specialized strategies of host manipulation. Molecular signatures of phagocytosis are present in all phytomyxean datasets analysed; but the model aggregates datasets in different trophic modes according to genome-based and transcriptome-based signatures (Fig. 1). These different predictions can be explained in the light of the polyphasic phytomyxean life-cycle (Liu *et al.*, 2020), where the transcriptome provides a realized molecular snapshot of the feeding stage whilst the genome also contains genetic information on stages other than the intracellular plasmodium (e.g., free-living flagellated zoospores). Genomic signatures identify Phytomyxea as phago-prototrophs whilst transcriptomes of intracellular parasitic stages are best explained by the subset of signatures of the intracellular phagotrophic specialist *Rozella allomycis* (Powell *et al.*, 2017; Fig. 1). Molecular signatures associated with the flagellum are the main drivers assigning *P. brassicae* genomes to the phago-prototrophic niche. Whilst flagella are associated with phagotrophy in certain organisms (e.g. Choanozoa); in Phytomyxea, flagella are exclusively associated with locomotion in the zoosporic phase of the life-cycle but find no implications in nutrition (Barr & Allan, 1982; Parodi *et al.*, 2010; Feng *et al.*, 2012). On the other hand, molecular signatures belonging to the Ras GTPases, mTORC1 and mTORC2 complexes are shared between *P. brassicae* and *M. ectocarpii* transcriptomes, both assigned to the *Rozella-like* phagotrophic specialists. Ras GTPases are known to control cytoskeletal remodelling and vesicular trafficking in human phagocytes (Wiedemann *et al.*, 2005) and mTORC2 has been linked to cytoskeletal polarization related to budding in yeasts (Loewith *et al.*, 2002). Furthermore, mTOR complexes, particularly mTORC1, are known to be paramount sensors of the nutritional state of the cell acting as a switch between anabolic and catabolic metabolism; and more broadly between growth and proliferation on one hand and autophagy and apoptosis on the other (Sabatini, 2017; Condon & Sabatini, 2019). This hints at a pivotal role of perception of the nutritional state and signal transduction in the intracellular feeding plasmodia of phytomyxids, coupled with cytoskeletal rearrangements that are paramount for phagocytic behaviour.

Molecular evidence shows that phytomyxean plasmodia rely on a reduced molecular machinery to perform intracellular phagotrophy, similarly to what happens in the intracellular fungal parasite *Rozella allomycis* (Burns *et al.*, 2018). *Rozella allomycis* belongs to the Cryptomycota, an early-diverged group within the true Fungi (James *et al.*, 2013) which has been demonstrated to be capable of intracellular phagocytosis (Powell *et al.*, 2017). Ultrastructural and molecular data agree in showing that *Rozella allomycis* mitochondria are non-functional and that the parasite relies on the host ones (James *et al.*, 2013; Powell *et al.*, 2017), thus hinting at a complete trophic reliance on the host. Despite the wide phylogenetic distance separating Cryptomycota and Phytomyxea, molecular signatures of phagotrophy from *Rozella* seem to accurately describe the behaviour of intracellular plasmodia in Phytomyxea.

Trophic dependency in intracellular plasmodia of Phytomyxea is also supported by ultrastructural observations of *P. brassicae* and *M. ectocarpii*, highlighting big, electron translucent and nearly featureless mitochondria (Supplementary Figure 4). In *R. allomycis* a similar mitochondrial morphology was linked to complete trophic dependency on the host in the intracellular stage (Powell *et al.*, 2017). The smaller and denser mitochondria with defined cristae observed in phytomyxean spores (Supplementary Figure 4; Talley *et al.*, 1978;) and the already reported co-presence of microbodies in zoospores (Tanaka *et al.*, 2001) is consistent with the usage of stored lipids as primary energy source (Held, 1975; Powell *et al.*, 2017). Taken altogether this evidence suggests a metabolic switch from complete reliance on the host during the intracellular growth, to zoosporic reliance on stored fatty acids in Phytomyxea and relates it to distinct changes in mitochondrial activity. Indeed, in contrast to *Rozella*, analyses of the mitochondrial genome of *P. brassicae* did not highlight any particular lack in functional genes (Daval *et al.*, 2018; Stjelja *et al.*, 2019), suggesting that the mitochondrion is still completely functional in other parts of the life cycle.

Phagocytosis in Phytomyxea has adapted to their intracellular lifestyle. Feeding plasmodia appear to uptake host structures by protrusion of lobes and invagination of the plasma membrane, again reminding processes observed in *R. allomycis* (Powell *et al.*, 2017), but also akin to the so-called “prey infiltration” strategy used by phylogenetically-related free-living amoebae in the order Vampyrellida (Hess & Suthaus, 2022). Our observations in TEM and fluorescent microscopy show different host organelles engulfed by *M. ectocarpii*, although phagocytosis seem skewed towards phaeoplasts (Fig. 2 and Fig. 4 E-H). Likewise, *P. brassicae* seems to target preferentially the host amyloplasts (Fig. 3 and Fig. 4 A-D). Whether this seemingly selective uptake of host organelles is the outcome of real targeting, of chance (phaeoplasts and amyloplasts are the most widespread organelles in the respective host cells) or an observational artefact caused by the delayed digestion of complex plastid-derived organelles remain to be ascertained.

However, it is known that host manipulation by *P. brassicae*, beside inducing mitosis and cellular expansion in the host (Olszak *et al.*, 2019); generates a strong physiological sink, driving photosynthates to the infected host cells (Malinowski *et al.*, 2019). Those photosynthates accumulate as starch in amyloplasts, which are significantly more abundant in the infected root cells of brassicas (Ma *et al.*, 2022) where they appear to be superficially “plugged” into the plasmodium surface, reminding of a process of semi-extracellular phagocytosis (named “pomacytosis”, Kamennaya *et al.*, 2018). Previous studies highlighted an upregulation of the plastidial MEX1 maltose transporter in infected roots, involved in the export of maltose outside the plastid after starch degradation (Badstöber *et al.*, 2020, Supplementary Figure 5). In this context, we can hypothesise that the pomacytosis-like process observed in *P. brassicae* co-opts phagocytosis to allow for the formation of a close interface between the parasite and the amyloplast without cutting the organelle away from the nucleus, thus allowing for the leaking and fast uptake of soluble sugars such as glucose.

Recently identified *P. brassicae* glucose transporters and glucose content in infected roots have been found to significantly increase in late stages of infection (Kong *et al.*, 2022). In the context of a biotrophic interaction that relies on phagotrophy, feeding specifically on amyloplasts has the clear advantage of targeting the host’s carbon storage, whilst at the same time avoiding organelles paramount to the host cell survival and regulation (e.g., the nucleus). This in turn, keeps intact the molecular machinery necessary for the host cell to continue accumulating photosynthetate as amyloplastic starch, giving time to the parasite to complete its life-cycle. Nonetheless, it is unclear whether the targeted phagocytosis/pomacytosis of amyloplasts is an active process or one passively driven by space-constraints within host cells packed with energy rich organelles (Fig. 3 and Fig. 4). Further evidence needs to be produced to confirm this hypothesis; but if confirmed, this would place *P. brassicae* in a particular ecological niche where ancestral phagocytosis provided the baseline to exploit host resources obtained via molecular manipulation co-evolved with specific hosts (Pérez-López *et al.*, 2020; Hossain *et al.*, 2021; Pérez-lópez *et al.*, 2021).

Results gathered from *M. ectocarpii* further support intracellular phagocytosis as main mode of nutrition within Phytomyxea. More so since, differently from *P. brassicae* data derived from sporogenic plasmodia, molecular and morphological data for *M. ectocarpii* come from sporangial plasmodia (Maier *et al.*, 2000). Following a conservative and parsimonious interpretation, our results therefore hint at a paramount role of intracellular phagocytosis in both the sporangial and sporogenic phases of the phytomyxean life cycle. *M. ectocarpii* also induces mitosis and cell expansion in its algal host (Maier et al., 2000) but evidence on carbohydrate accumulation in infected tissue has not yet been produced. Brown algae accumulates photosynthates mainly as soluble vacuolar laminaran and cytoplasmic mannitol (Michel *et al.*, 2010; Chabi *et al.*, 2021). It is therefore interesting to notice the early disappearance of the vacuole in cell infected by *M. ectocarpii.* Although consumption of the vacuole seems to be a necessary step of intracellular colonization, simply to provide growth space for the enlarging sporangium; this would also allow for the parasite to immediately consume the major polysaccharide storage within the host cell providing it with rapid energy. However, a first glance at levels of gene expression in Maullinia-infected *E. siliculosus* Ec32m did not highlight a clear pattern of upregulation of laminarin / mannitol catabolism or extra-vacuolar transport of carbohydrates (Supplementary Table 2), thus whether *M. ectocarpii* manipulates its host cell carbohydrate metabolism remains unclear. It is worth reminding that the transcriptome analysed here originated from an asynchronous parasite population, where the signal of a possibly transient and punctual interaction between the parasite and the host vacuole could have been diluted in the bulk approach used.

In the case of *M. ectocarpii*, phaeoplasts seemed to be a preferred target of phagocytosis. In fact, TEM images show that phaeoplasts in infected cells shrink with the progression of the infection (Fig.2 and Supplementary Figure 3). In depth investigation of plastidial dynamics in infected algal cells is beyond the scope of this study, but the possibility that plastids are directly manipulated before being targeted for consumption by the parasite is possible. If proven, this would hint at a conservation or convergence of the target host organelle within the Phytomyxida. Similar patterns of plastidial shrinkage have been highlighted in the interaction between the intracellular oomycete parasite *Anisolpidium ectocarpii* (infecting *M. pyrifera*), but in this case the decrease in size was interpreted as result of autophagic processes and thus to the reaction of the host against the parasite (Murúa *et al.*, 2020).

It is worth bearing in mind that only scarce information is available on the sporogenic stage of *M. ectocarpii* (Parodi *et al.*, 2010; Blake *et al.*, 2017) and on Phagomyxida overall (Schnepf & Bulman, 2000; Murúa *et al.*, 2017). If, as it is suspected, a sporogenic phase inducing galls formation in adult kelp sporophytes does exist (Blake *et al.*, 2017), an even higher degree of host manipulation can be expected for *M. ectocarpii*, bringing it even closer to its land-dwelling relative *P. brassicae*.

Although not common, intracellular endocytosis has been documented in intracellular parasites spanning the taxa Apicomplexa (Spielmann *et al.*, 2020 and references therein), Cryptomycota (Torruella *et al.*, 2018) and Euglenozoa (Etheridge, 2022). The reduced dataset of phagotrophy-related proteins from *Rozella allomycis* correctly describes *Trypanosoma cruzii* and *Leishmania braziliensis* as capable of intracellular phagotrophy (Chasen *et al.*, 2020; Halliday *et al.*, 2020); whilst fails to assign *Plasmodium falciparum* and *Toxoplasma gondii* to this category. Indeed, *P. falciparum* and *T. gondii* are known to use a different set of genes to undertake endocytic nutrients uptake (Spielmann *et al.*, 2020) and especially to lack important genes involved in small GTPase (RAS superfamily) and TOR signalling pathways (Van Dam *et al.*, 2011), which are paramount for the predictive model (Burns *et al.*, 2018). The apparent proximity of the genetic make-up underpinning intracellular phagocytosis in unrelated Phytomyxea, *Rozella* and trypanosomatids is intriguing, since it hints at the possibility that the smallest subset of genes required for phagocytosis is present in these otherwise unrelated parasites. Although speculative, this hypothesis suggests that intracellular parasites maintaining a phagocytic behaviour despite the well-known process of genome reduction (Keeling & Slamovits, 2005) might, when not overly specialised toward their host, make a good model to investigate the very core of the phagocytic machinery.

The data presented and discussed here place phytomyxean intracellular parasites half-way between the extremes of specialised biotrophic host manipulation and osmotrophy and generalist phagocytic predation. Growing molecular and microscopic evidence suggests that phagocytosis is a backbone feature of Rhizarians upon which “variations on the theme” brought about the diversification of the group (Anderson, 1978; Hirakawa, 2017; Gerbracht *et al.*, 2022; Hess & Suthaus, 2022). In this context, Phytomyxean are not an exception. It is tempting to speculate that the maintenance and adaptation of phagocytic behaviour is one of the reasons behind the success of this impactful and recalcitrant parasites; allowing them to specialise to certain hosts meanwhile maintaining the ability to feed and propagate within a broader set of organisms (Ludwig-Müller *et al.*, 1999; Maier *et al.*, 2000; Qu & Christ, 2006). Further research on this group of intriguing parasites will surely provide more evidence on the degree of host manipulation/phagocytosis within the class, especially if targeted towards non-model organisms for which data are lacking. Comparative investigations and the exploration of biodiversity surrounding parasites and pathogens proves paramount to deeply understand their biology and potentially devise strategies to counter their effects and broadly foresee the evolutionary trajectories of parasitism.

## Supporting information

**Supplementary figure 1**

Caption: Results of BUSCO analysis of the genomes and transcriptomes of all Phytomyxid analysed supporting the high completeness.

**Supplementary figure 2**

Caption: Optical and fluorescence micrographs provide evidence of phagocytosis in intracellular plasmodia of *Maullinia ectocarpii* in *Macrocystis pyrifera* (A-G). Images have been captured using differential interference contrast microscopy (DIC; A”-G”) and fluorescence microscopy (A’-G’) and subsequently overlaid (A-G). FISH-probes specific to the 18S rRNA gene of *M. ectocarpii* were used to highlight the ribosome rich cytoplasm of the parasites (green). Hoechst staining highlighted the nuclei of both parasite and host (blue) in fluorescence microscopy. The white arrow in B and F points toward the Hoechst-stained host cell nucleus, whilst white arrowheads in A, B, C, and F indicate putative phagocytic vacuoles. Those are further identified by numbers in A, B and C and highlighted by a white border in F. No algal organelles are clearly discernible within the digestive vacuoles in A-D. Besides absence, this might also be due to the overall coarse appearance of the parasitic thallus and/or to the tilted position of the investigated cell. The plasmodium in E-G is at a later stage of development as highlighted by the complete absence of visible algal organelles (especially phaeoplasts) with the exception of the phagocytised nucleus. Focal planes represent a high (A, A’, A” and D, D’, D”), a central (B, B’, B”; E, E’, E” and F, F’, F”) and a low (C, C, C” and G, G’, G”) layer from z-stacks (steps = 0,1 μm) containing entire plasmodia. All scale bars are 10 μm.

**Supplementary Figure 3**

Caption: The picture features a young plasmodium of *Maullinia ectocarpii* in its host alga *Macrocystis pyrifera.* The young age of the plasmodium can be inferred from the presence of only one nucleus. The plasmodium (highlighted by a black line in A’) bears 4 pseudopodia-like structures (Fig. A’, Ps 1-4), 2 of which (Fig. A’, Ps3 and Ps4) are used to surround and engulf an algal organelle of difficult identification (Fig A’, ?). List of abbreviations : **N** = parasite nucleus, **HMt** = host mitochondria, **HPl** = host phaeoplasts, **CV** = central algal vacuole, **asterisks (*)** = potential parasite mitochondria, **black lines** indicate the plasma membrane of the plasmodium, **Ps** = pseudopodia. Scale bars = 500 nm.

**Supplementary Figure 4**

Caption: Comparison of mitochondria between *Maullinia ectocarpii* (left) and *Plasmodiophora brassicae* (right); between feeding plasmodia (top) and spores (bottom). Plasmodial mitochondria are bigger, have fewer, less defined cristae and a sparse electron translucent matrix; compared with the electron-dense and well-defined mitochondria encountered in spores. Note: the figure shows flagellated secondary zoospores for *M. ectocarpii*, whilst thick-walled resting spores are shown for *P. brassicae*. Scale bars 200 nm.

**Supplementary Figure 5**

Caption: Changes in the host plant *Brassica oleracea* var. *gongylodes* starch metabolism during infection from *Plasmodiophora brassicae* as per Ciaghi et al., 2018. Log 2 fold changes for significantly differentially expressed genes are shown in the heatmap, which includes numerical transcript identifiers and annotation according to MapMan/Mercator. Up-regulated genes are shaded red, whilst down-regulated genes are shaded blue. The yellow/brown figure symbolizes the amyloplast. The maltose transporter MEX1 and its upregulation in both white and brown root galls as compared to symptomless roots is further highlighted by the white box on the amyloplast wall. Naming conventions follow Ciaghi et al., 2018: WG = small white spindle galls, BG = larger brownish spindle galls and SL = symptomless roots. For details about biological material, data and analyses please refer to Ciaghi et al., 2018.

**Supplementary Material 1**

Sheet 1 - Caption: Comparison between best-hit genes against molecular signatures of general phagotrophy between P.brassicae e3 genome and transcriptome.

Sheet 2 - Caption: Comparison between best-hit genes against molecular signatures of Rozella-like specialist phagotrophy between P.brassicae and M.ectocarpii transcriptomes.

**Supplementary Material 2**

Video – from Fig. 2

Because of file size issues, videos from Fig.2 have been made available via the following link to the file sharing service of the University of Innsbruck.

https://fileshare.uibk.ac.at/d/c729cb8ceb6446c1a0ed/

**Supplementary Material 3**

Video – from Supplementary Figure 2

Because of file size issues, videos from Supplementary Figure 2 have been made available via the following link to the file sharing service of the University of Innsbruck.

https://fileshare.uibk.ac.at/d/8799738424d0462b8fae/

**Supplementary Table 1**

Caption: Values from the TrophicModePredictionTool assigning the proteomes of the listed organisms to 5 different trophic modes. Predictions are deemed significantly accurate when above 0.5 and increase in accuracy with higher values.

**Supplementary Table 2**

Caption: Log 2 fold change expression levels in *Maullinia*-infected *E. siliculosus* Ec32m versus non-infected control for enzymes involved in carbohydrate metabolism. The list of genes presented was compiled based on data from Michel *et al.*, 2010; Chabi *et al.*, 2021 and further manually curated.

