## Supplementary Figure for "Phagocytosis underpins the biotrophic lifestyle of intracellular parasites in the class Phytomyxea (Rhizaria)"

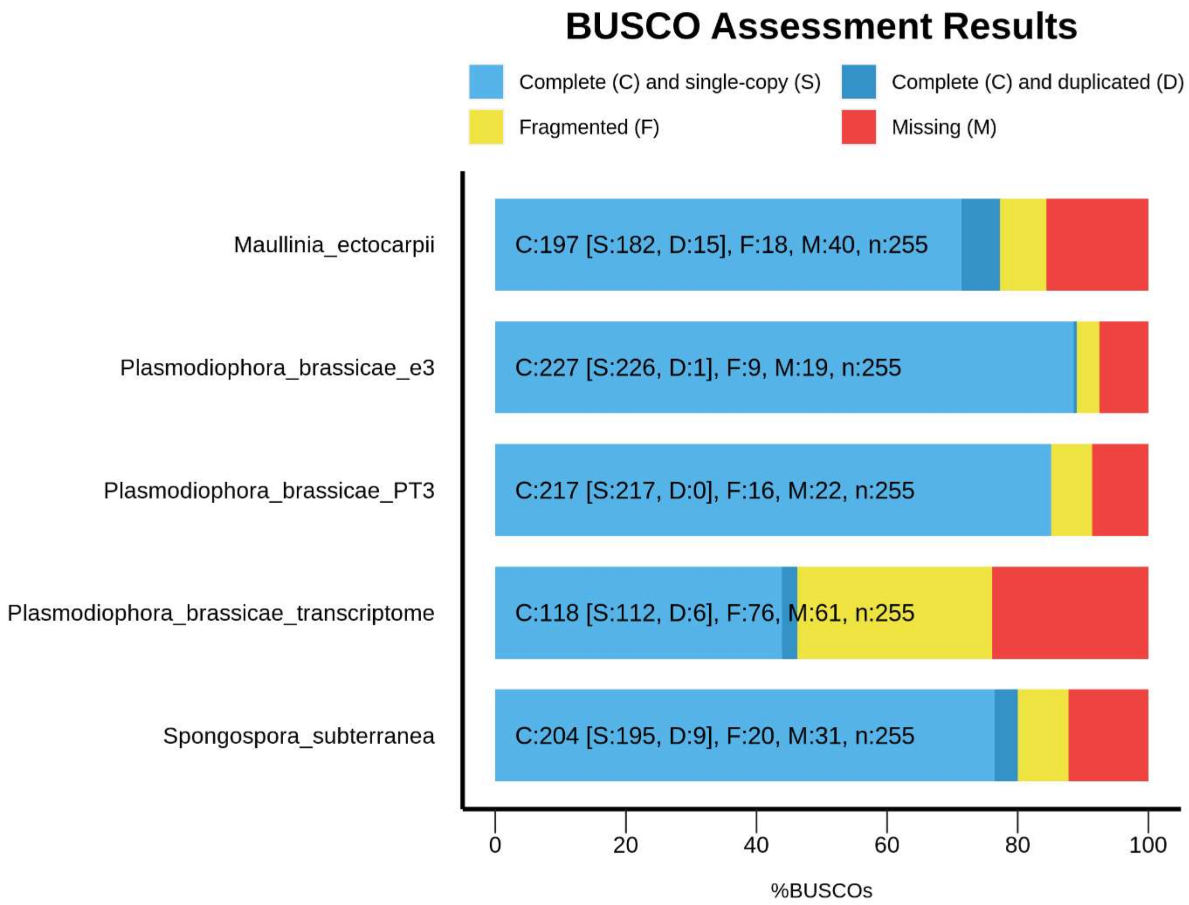


**Supplementary figure 1**

Caption: Results of BUSCO analysis of the genomes and transcriptomes of all Phytomyxid analysed supporting the high completeness.

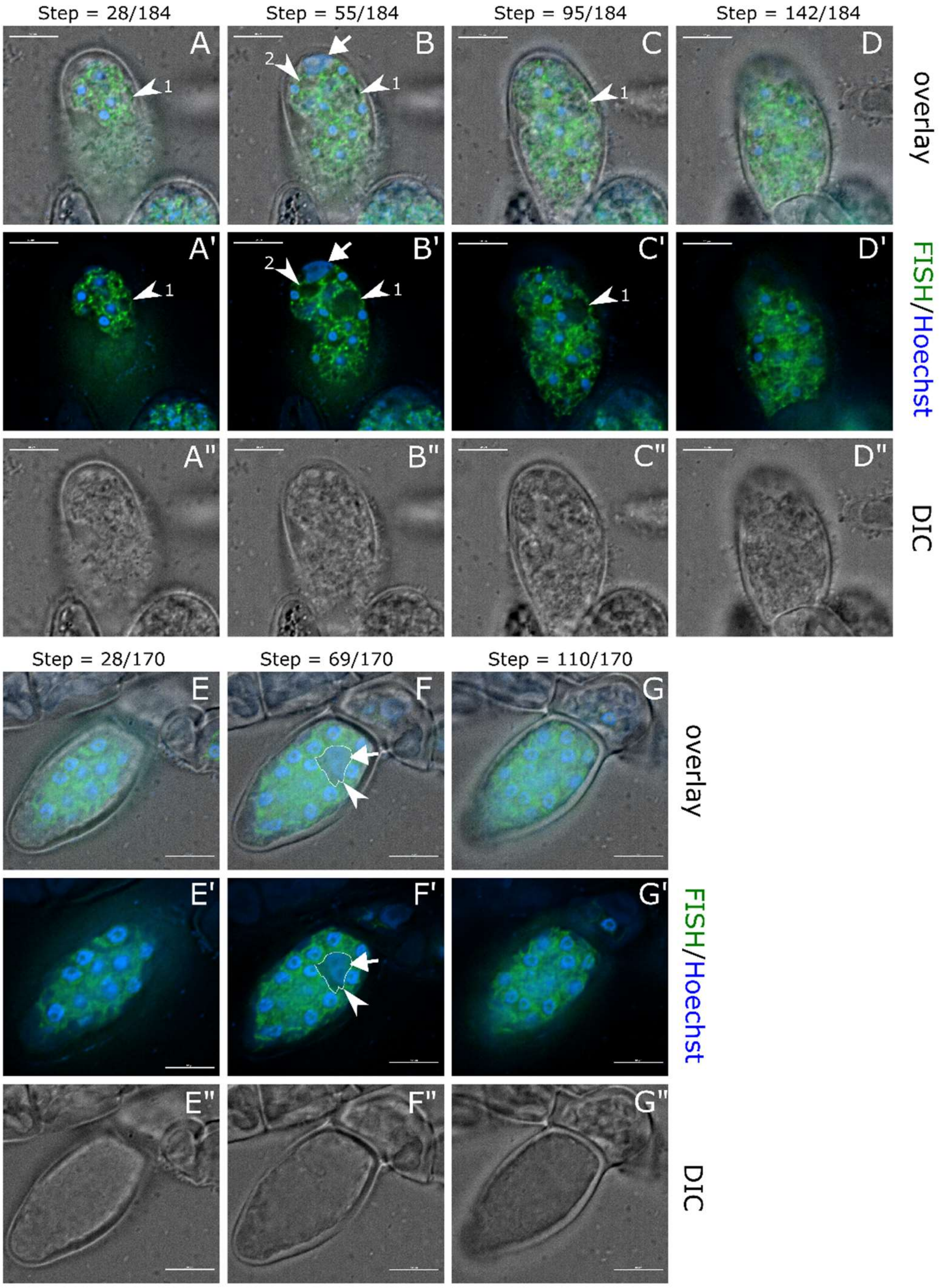


**Supplementary figure 2**

Caption: Optical and fluorescence micrographs provide evidence of phagocytosis in intracellular plasmodia of *Maullinia ectocarpii* in *Macrocystis pyrifera* (A-G). Images have been captured using differential interference contrast microscopy (DIC; A"-G") and fluorescence microscopy (A'-G') and subsequently overlaid (A-G). FISH-probes specific to the 18S rRNA gene of *M. ectocarpii* were used to highlight the ribosome rich cytoplasm of the parasites (green). Hoechst staining highlighted the nuclei of both parasite and host (blue) in fluorescence microscopy. The white arrow in B and F points toward the Hoechst-stained host cell nucleus, whilst white arrowheads in A, B, C, and F indicate putative phagocytic vacuoles. Those are further identified by numbers in A, B and C and highlighted by a white border in F. No algal organelles are clearly discernible within the digestive vacuoles in A-D. Besides absence, this might also be due to the overall coarse appearance of the parasitic thallus and/or to the tilted position of the investigated cell. The plasmodium in E-G is at a later stage of development as highlighted by the complete absence of visible algal organelles (especially phaeoplasts) with the exception of the phagocytised nucleus. Focal planes represent a high (A, A', A" and D, D', D"), a central (B, B', B"; E, E', E" and F, F', F") and a low (C, C, C" and G, G', G") layer from z-stacks (steps = 0,1 µm) containing entire plasmodia. All scale bars are 10 µm.

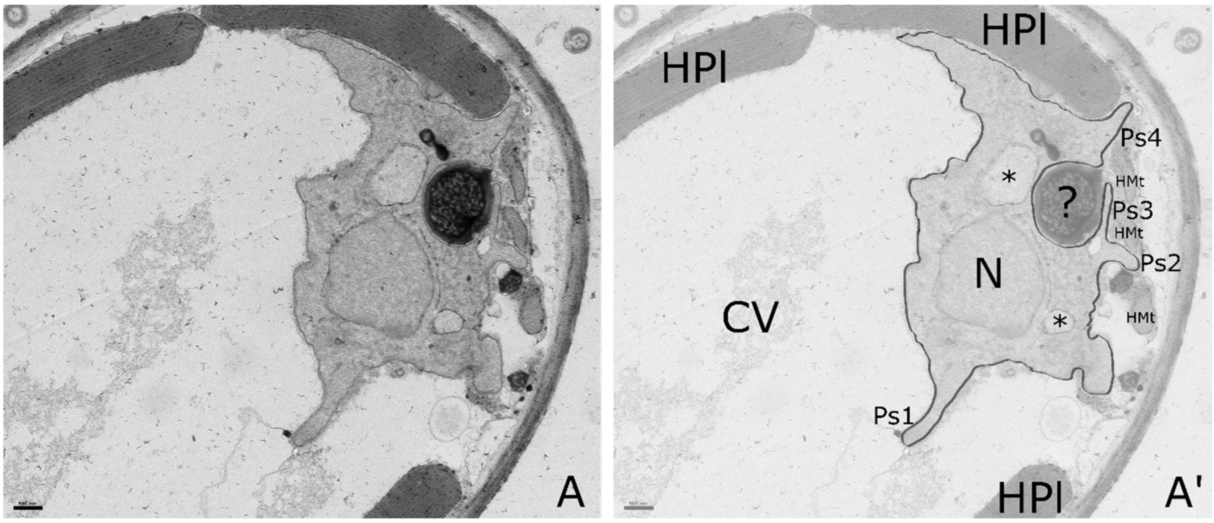


**Supplementary Figure 3**

Caption: The picture features a young plasmodium of *Maullinia ectocarpii* in its host alga *Macrocystis pyrifera*. The young age of the plasmodium can be inferred from the presence of only one nucleus. The plasmodium (highlighted by a black line in A') bears 4 pseudopodia-like structures (Fig. A', Ps 1-4), 2 of which (Fig. A', Ps3 and Ps4) are used to surround and engulf an algal organelle of difficult identification (Fig A', ?). List of abbreviations : **N** = parasite nucleus, **HMt** = host mitochondria, **HPl** = host phaeoplasts, **CV** = central algal vacuole, **asterisks (*)** = potential parasite mitochondria, **black lines** indicate the plasma membrane of the plasmodium, **Ps** = pseudopodia. Scale bars = 500 nm.

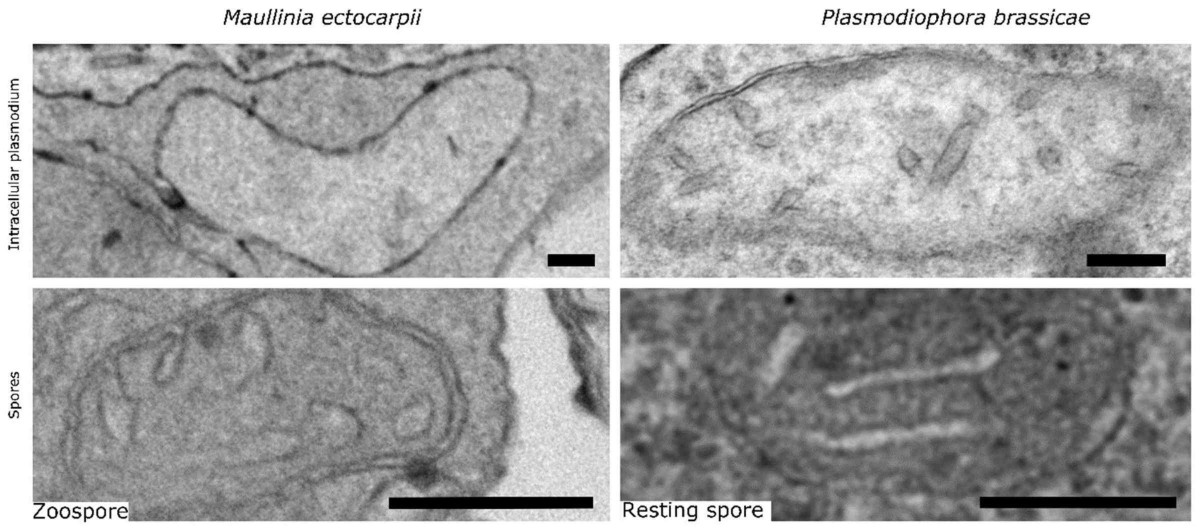


**Supplementary Figure 4**

Caption: Comparison of mitochondria between *Maullinia ectocarpii* (left) and *Plasmodiophora brassicae* (right); between feeding plasmodia (top) and spores (bottom). Plasmodial mitochondria are bigger, have fewer, less defined cristae and a sparse electron translucent matrix; compared with the electron-dense and well-defined mitochondria encountered in spores. Note: the figure shows flagellated secondary zoospores for *M. ectocarpii*, whilst thick-walled resting spores are shown for *P. brassicae*. Scale bars 200 nm.

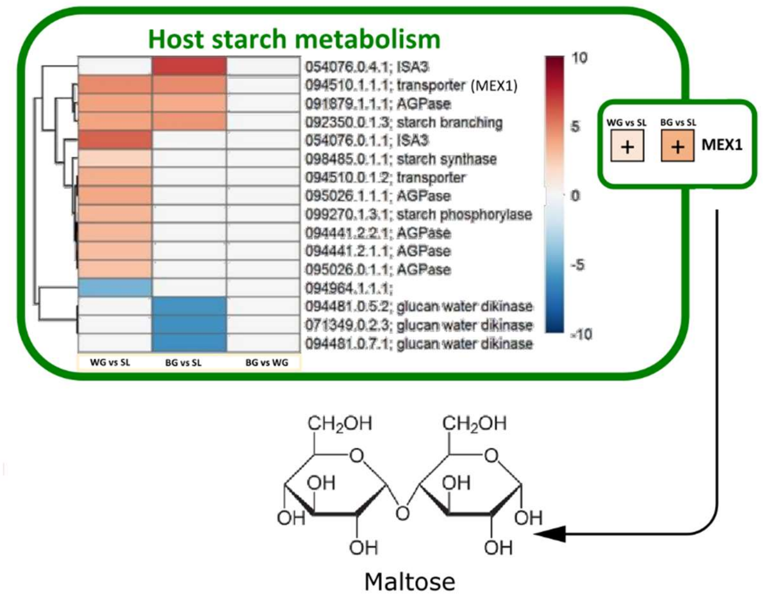


Caption: Changes in the host plant *Brassica oleracea* var. *gongylodes* starch metabolism during infection from *Plasmodiophora brassicae* as per Ciaghi et al., 2018. Log 2 fold changes for significantly differentially expressed genes are shown in the heatmap, which includes numerical transcript identifiers and annotation according to MapMan/Mercator. Up-regulated genes are shaded red, whilst down-regulated genes are shaded blue. The yellow/brown figure symbolizes the amyloplast. The maltose transporter MEX1 and its upregulation in both white and brown root galls as compared to symptomless roots is further highlighted by the white box on the amyloplast wall. Naming conventions follow Ciaghi et al., 2018: WG = small white spindle galls, BG = larger brownish spindle galls and SL = symptomless roots. For details about biological material, data and analyses please refer to Ciaghi et al., 2018.
